# Developmental changes in diffusion markers of neurite vary across the hippocampus and covary with the cellular composition of hippocampal tissue

**DOI:** 10.1101/2024.09.04.611231

**Authors:** J. Kember, Z. Gracia-Tabuenca, R. Patel, M. Chakravarty, X.J. Chai

## Abstract

The hippocampus is a critical brain structure supporting memory encoding and retrieval, yet the development of its microstructure in humans remains unknown. Understanding this development may provide insight into the mechanisms underlying memory and their disruption in disease. To address this, we non-invasively estimated the density and branching complexity of neurite (dendrites, axons, glial processes) using diffusion-weighted MRI in 364 participants aged 8–21. With development, we observed large increases in neurite density and branching complexity that persisted until approximately 15 years of age before stabilizing at adult-like values. Increases in neurite density were relatively homogenous across hippocampal axes, whereas increases in branching complexity were heterogeneous; increasing primarily in CA1, SRLM, subiculum, and anterior hippocampus. To assess whether this development may be attributable to specific cell-types, we tested for spatial overlap between age-related change in neurite and the cell-type composition of hippocampal tissue via cross-reference with an out-of-sample gene-expression atlas. We found age-related changes in neurite density spatially overlapped with a granule cell component; whereas age-related changes in neurite branching complexity overlapped with a pyramidal neuron component. These results provide the first glimpse at the nonlinear maturation of hippocampal microstructure and the cell-type composition of hippocampal tissue underlying these changes.

## Introduction

In the hippocampus, thin extensions from the cell soma– axons, dendrites, glial processes– collectively known as neurites, play a fundamental role in shaping circuits and in the emergence of functional processes. Histological and quantitative-MRI findings indicate the morphological properties of neurites, including their density and branching complexity, vary systematically along the transverse and longitudinal axes of the hippocampus (Beaujoin et al., 2018; Karat et al., 2023). This spatial variation has important implications for function, as a close relationship exists between the morphological properties of neurites and the computational properties of both neurons (e.g., membrane capacitance), and circuits (e.g., the generation and modulation of neurophysiological oscillations; Andrade-Talavera et al., 2023; Kota et al., 2020; Malik et al., 2016; Shafiei et al., 2023; Stuart et al., 2015; Tukker et al., 2013; Vogel et al., 2021).

Late into adolescent development, the density and branching complexity of neurites continues to increase across the association cortices, unfolding in a regionally specific manner which aligns closely with the sensorimotor–association cortex hierarchy (Lynch et al., 2023; Sydnor et al., 2021). Indeed, while the density and branching complexity of sensory cortices shows little to no change following early childhood, neurite density of dorsal medial prefrontal cortex, and branching complexity of dorsolateral prefrontal/midcingulate cortex continues to increase beyond 20 years of age (Lynch et al., 2023; Zhao et al., 2021). These findings are in line with work in rats, where the length and spine-density of both basal and apical dendrites in the PFC peaks in adolescence (Juraska & Willing, 2017). These maturational changes have been partially attributed to pubertal increases in dehydroepiandrosterone (DHEA; Juraska & Willingl, 2017).

Despite this substantial maturation in neurite morphology observed across the neocortex, as well as the functional relevance of neurites in the hippocampus, little is known about the development of neurite morphology within the hippocampus specifically. Understanding this development may be crucial in explaining the basic mechanisms underlying hippocampal function (Ofen et al., 2019), as well as their disruption in certain neurodevelopmental disorders (i.e., ASD; Li et al., 2019).

So far the only study to examine developmental changes in neurite density and branching complexity in the hippocampus found increases in neurite density, and little to no change in branching complexity, from 8 to 14 years of age (Mah et al., 2017). However, this work averaged neurite estimates across the entire hippocampus, ignoring the prominent differences in microstructure that exist across both the medial and longitudinal axes. The primary dimension of this microstructural organization is across subfields: distinct regions topographically localized across the medial-lateral axis which include CA1–CA4, dentate gyrus, and the subiculum (Andersen et al., 2007). The second most prominent dimension of microstructural organization is along the anterior to posterior longitudinal axis of the hippocampus (Masouleh et al., 2020). Given the distinct structural and functional properties (Karat et al., 2023) and the differential development of these subregions (Vinci-Booher et al., 2023), we hypothesized that microstructural development would vary among the subfields and along the longitudinal axis. The defining microstructural properties of each subfield are typically identified from post-mortem histological tissue samples. However, recent work with diffusion-weighted magnetic resonance imaging (MRI) suggests that microstructural properties of hippocampal substructures can be delineated non-invasively and *in vivo* (Beaujoin et al., 2018; Karat et al., 2023; Zhang et al., 2012).

In the current study, we sought to identify age-related differences in the properties of neurites across the primary axes of hippocampal organization between childhood and young adulthood. To do so, we examined how diffusion-MRI markers of neurite density and branching complexity differed across subfields and along the longitudinal axis in 364 participants between 8 and 21 years of age from the Lifespan Human Connectome ProjectDevelopment (HCP-D) data, using Neurite Orientation Dispersion and Density Imaging (NODDI). Then, to understand the cellular composition of hippocampal tissue which exhibits these maturational changes in neurite, we investigated the spatial correlation between the cellular composition of hippocampal tissue and age-related changes in neurite morphology. We did so using an imaging transcriptomics approach, where we spatially integrated our neuroimaging data with transcriptomic data from the Allen human brain gene-expression atlas, while utilizing cell-type specific gene expression signatures identified from single-nuclei RNA sequencing of the human hippocampus (Arnatkevic□iūtė et al., 2019; Ayhan et al., 2021; Hawrylycz et al., 2012; Paus et al., 2023).

## Results

### Differences in neurite properties across hippocampal subfields and the longitudinal axis

To quantify neurite density and branching complexity, we examined the neurite density and orientation dispersion indices derived from NODDI, respectively (Neurite density Index: NDI, Orientation dispersion index: ODI; Zhang et al., 2012). We extracted these indices in 5 hippocampal subfields (CA1, SRLM, DG/CA4, CA3/CA2, Subiculum; average across right/left hippocampus) from each subject’s diffusion MRI imaging data (see Fig. 1a; see Methods for subfield segmentation). For longitudinal-position analyses, we divided each subject’s hippocampal mask into three parts of equal length along the longitudinal axis (see Fig. 1b). We then derived the average of each microstructural metric across each of these three segments: the anterior (hippocampus head), middle (body), and posterior third (tail), similar to published approaches (Solar et al., 2021; DeMaster et al., 2014; Lee et al., 2020).

**Fig 1.**
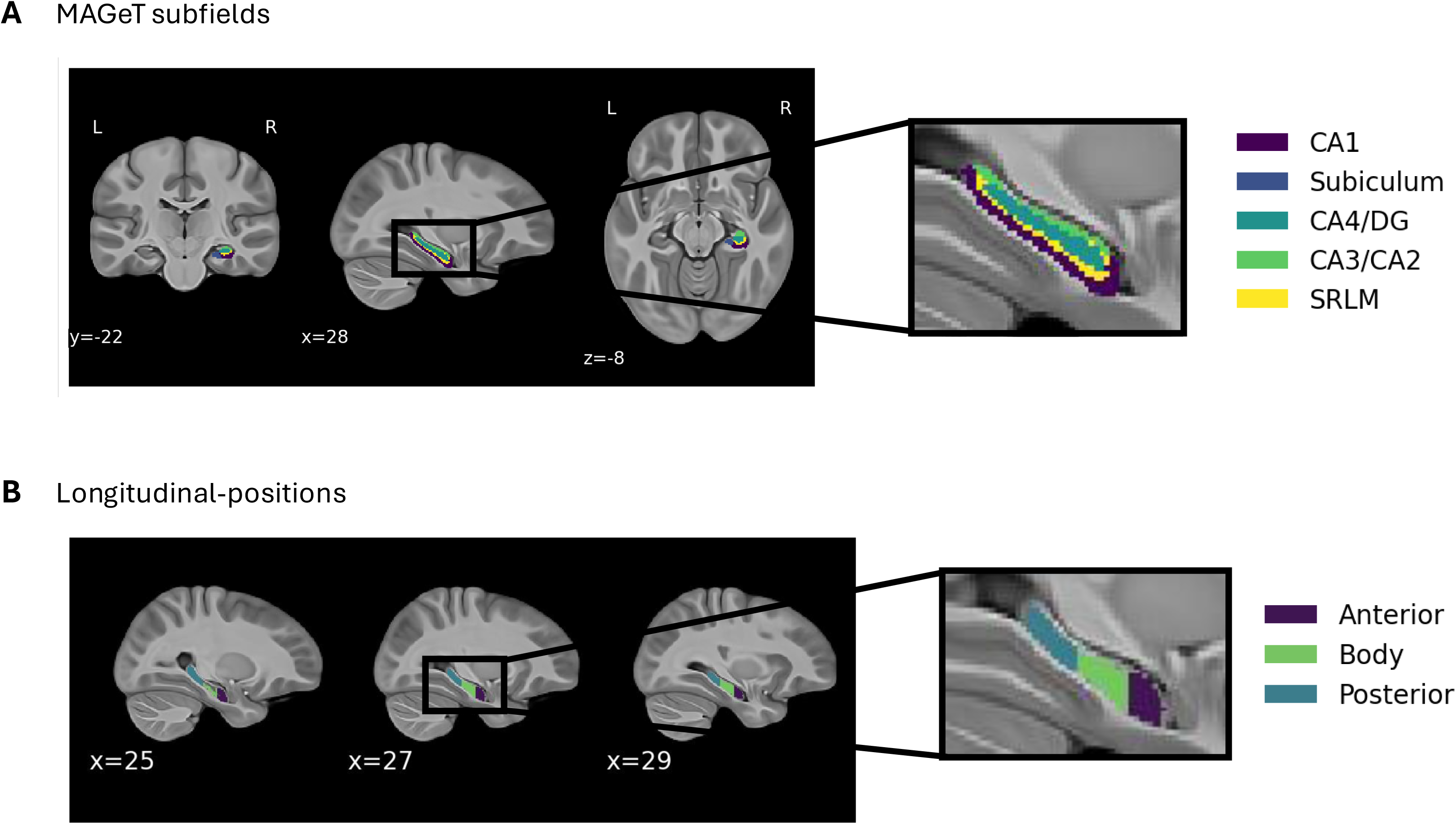
Hippocampal subfields and longitudinal-positions. (A) Group-average (majority-vote) subfields, derived from the MAGeT segmentations, overlaid on a template MNI-152 brain (DG = Dentate gyrus, CA =. Cornu ammonis, SRLM: stratum lacunosum moleculare). (B) Anterior, body, and posterior division of the hippocampus along the longitudinal axis.

Box-plots illustrating the density and branching complexity of neurite in each subfield are shown in Fig. 2a. For neurite density [NDI], estimates were highest in the subiculum and lowest in SRLM and CA4/DG. Subfield-by-subfield *t*-tests revealed that all subfields differed significantly from one another (FDR-*ps* < .05). For branching complexity [ODI], estimates were highest in SRLM and CA1, and lowest in CA3/CA2. All subfields differed significantly from one another (FDR-*ps* < .05). Bivariate subfield-by-subfield effect sizes (Cohen’s *d*) are shown in supplementary Fig. 1a; both NDI and ODI showed a range of small to very-large effect size differences across subfields.

**Fig 2.**
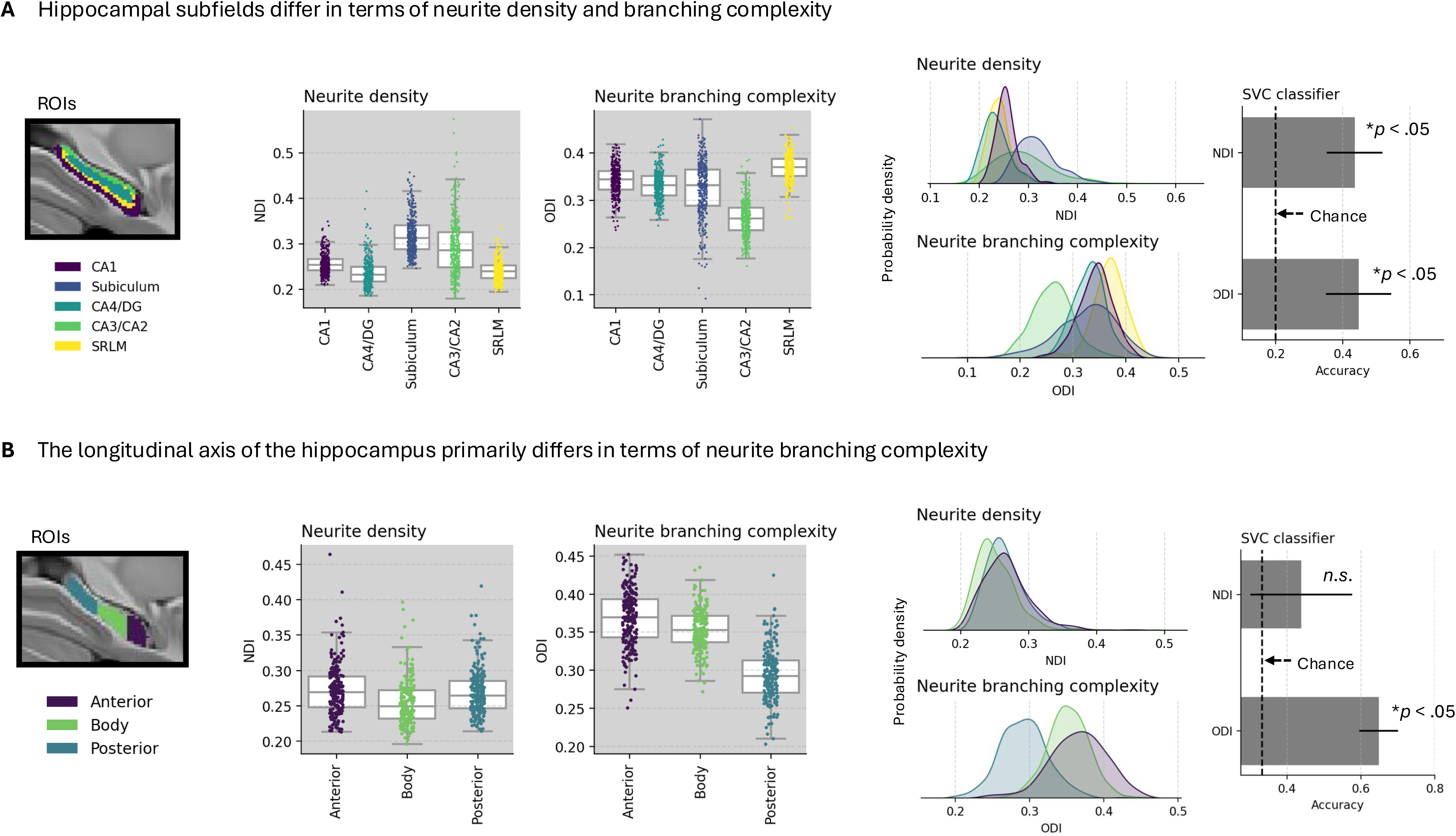
Differences in neurite density and branching complexity across the hippocampus. (A) On the left, box-plots illustrate the average neurite density and branching complexity across hippocampal subfields for each subject. On the right, the accuracies of support-vector classifiers trained to distinguish subfields based on either NDI or ODI, along with kernel-density estimates that allow for visualization of this differentiability across subfields. Chance-level accuracy of the classifiers is .20 (for 5 subfields). (B) Same as panel *A*, for longitudinal positions. Chance-level accuracy of the classifiers is .33 (for 3 longitudinal positions).

We then tested whether subfields primarily differ in terms of neurite density (NDI) or branching complexity (ODI). To do so, we compared the ability of separate machine learning models (support vector classifiers) to classify subfields based on either NDI or ODI. Accuracies of both the model trained on NDI and the model trained on ODI were above chance-level [see Fig. 2a; NDI: 43.5%, ODI: 44.7%, chance: 20%]. However, accuracy of the NDI and ODI models were not significantly different from one another, suggesting subfields differ in terms of NDI and ODI relatively equally.

We then conducted similar analyses for the longitudinal axis. Box-plots illustrating the density and branching complexity of neurite across the longitudinal axis are shown in Fig. 2b. For neurite density (NDI), estimates did not differ between anterior and posterior hippocampus, although both anterior and posterior showed significantly greater NDI than the hippocampal body (Anterior > Body: *t*_363_ = 9.01, *p* < 1e-15; Posterior > Body: *t*_363_ = 17.8, *p* < 1e-50). For branching complexity (ODI), a gradient along the longitudinal axis was revealed, whereby estimates were highest in the anterior hippocampus, followed by the hippocampal body, followed by the posterior hippocampus. All longitudinal positions differed significantly from one another (FDR-*ps* < .05). Bivariate position-by-position effect sizes (Cohen’s *d*) are shown in supplementary Fig. 1b; both NDI and ODI showed a range of small to very-large effect size differences across longitudinal positions (*d’*s ranging from .41 to 2.07). This gradient in ODI was apparent when inspecting values at a millimeter resolution along the longitudinal axis, suggesting they are not sensitive to our specific choice of parcellation (see supplementary Figure 4b).

We then tested whether subregions along the longitudinal axis primarily differ in terms of neurite density [NDI] or branching complexity [ODI] through the support vector classifier approach described above. Accuracy of the model trained on ODI was significantly higher than accuracy of the model trained on NDI (which was not above chance-level; see Fig. 2b). This suggests positions along the longitudinal axis differ primarily in terms of neurite branching complexity, as opposed to neurite density.

### Age-related differences in neurite properties across subfields and longitudinal position

We then examined how neurite density and branching complexity relate with differences in age by correlating NDI and ODI with age (Pearson’s *r*) at each hippocampal ROI. All subfields showed large effect-size increases in neurite density [NDI] with age (*r’s* ≥ .55, *p’s* < 1e-30). Age-related differences in neurite branching complexity [ODI], however, were more heterogeneous across subfields: large effect-size increases were observed in CA1, Subiculum, and SRLM (*r*’s ≥ .43, *p’s* < 1e-16), a small effect-size increase in ODI was observed in CA4/DG (*r* = .11, *p* = .03), and a trend towards decreased ODI was observed in CA3/CA2 (*r* = -.09, *p* = .06). Prominent increases in both neurite density and branching complexity were observed along the entire longitudinal axis, although the strength of these effects varied by position: larger effect-size increases in neurite density [NDI] were observed in the body and posterior hippocampus compared to the anterior hippocampus. Whereas larger effect-size increases in neurite branching complexity [ODI] were observed in the anterior hippocampus compared to the body and posterior hippocampus. The strength of all correlations with age, as well as 95% confidence intervals, are shown in Fig. 3.

**Fig 3.**
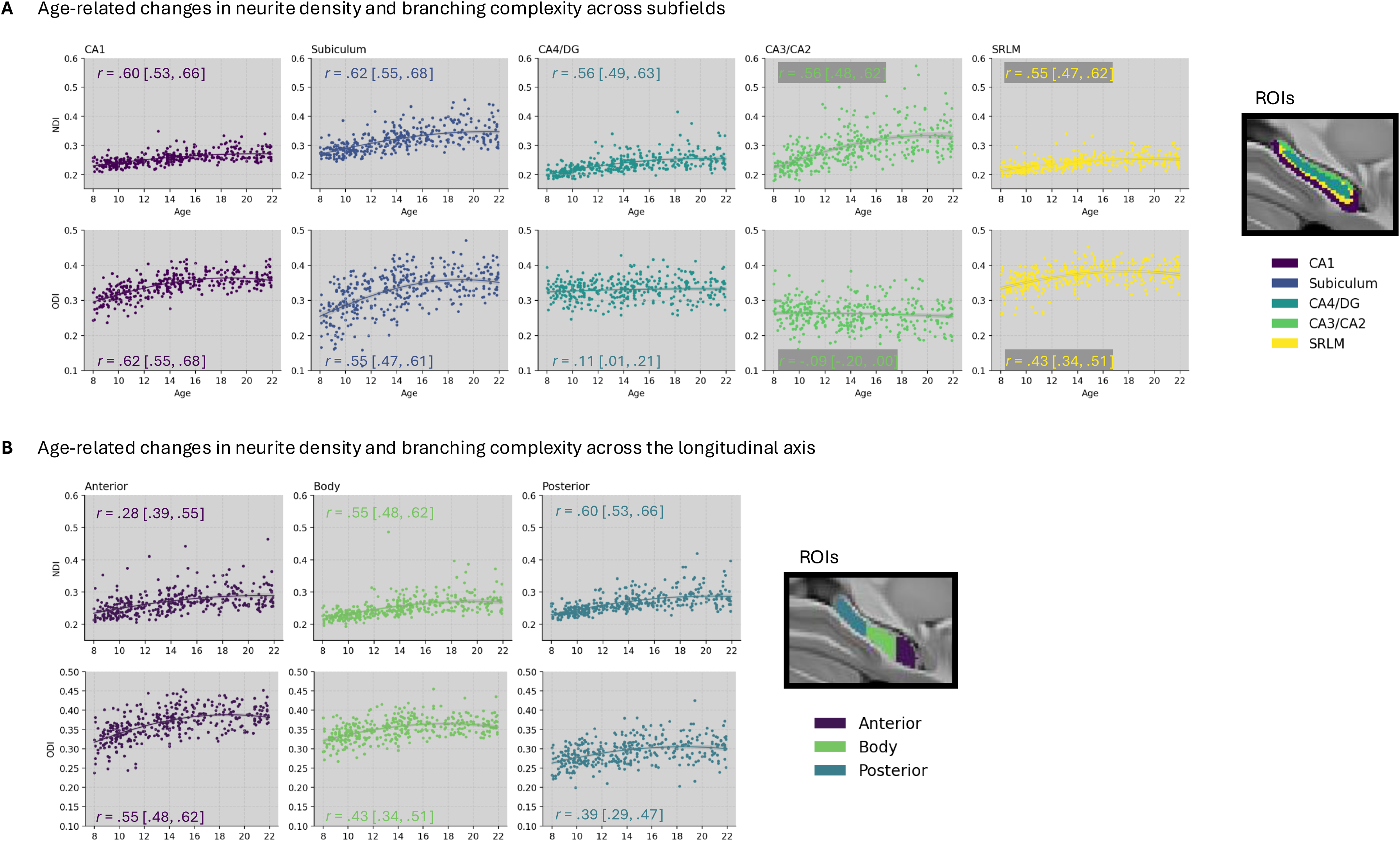
Development of neurite density and branching complexity across the hippocampus. (A) Scatter-plots illustrating the correlation between age and individual differences in neurite density (NDI: top panel) and neurite branching complexity (ODI: bottom panel) for each hippocampal subfield. Non-linear trajectories, along with shaded 95% confidence intervals are overlaid on scatter-plots. The correlation with age (Pearson’s *r*) and 95% confidence intervals are presented on each scatter-plot. (B) Same as panel *A*, for longitudinal positions.

Next, to capture any nonlinear changes that may occur with development, we estimated neurite properties as non-linear functions of age using a locally-weighted regression, with smoothing estimated through cross-validation (see methods). Visual inspection of the developmental trajectories of neurite density [NDI] revealed a common pattern across the hippocampus: strong increases in childhood that tapered off during adolescence. This general pattern, which can be seen in Fig. 3, was observed for each subfield and all longitudinal positions, and mirrors changes in neurite density observed across higher-order association cortex (Lynch et al., 2023). For neurite branching complexity [ODI], a similar pattern was observed for CA1, Subiculum, and SRLM, and across longitudinal positions. By contrast, CA4/DG and CA3/CA2 showed little to no change with age. To quantify these patterns, we estimated the age at which values stopped differing significantly from adults (subjects aged 18 and older, whose subregion microstructures can be seen in supplementary Figure 2), putatively reflecting time-points in development where maturational processes stop. These ages can be seen in Fig. 4. Across subfields, changes in neurite density [NDI] became not-significantly different from adults at roughly 16 years of age, with little variation across subfields. By contrast, changes in neurite branching complexity differed considerably across subfields: there was no significant difference with adults at any age for CA3/CA2 or CA4/DG; no difference following 14.4 years of age in CA1 and subiculum; and no change following 12.7 years of age in SRLM. Across the longitudinal axis, neurite density stopped differing from adults at 15.1 years, 15.9 years, and 17.0 years for the anterior, body, and posterior hippocampus, respectively. Whereas neurite branching complexity stopped differing from adults at 14.5 years, 13.6 years, and 13.8 years for the anterior, body, and posterior hippocampus, respectively.

**Fig 4.**
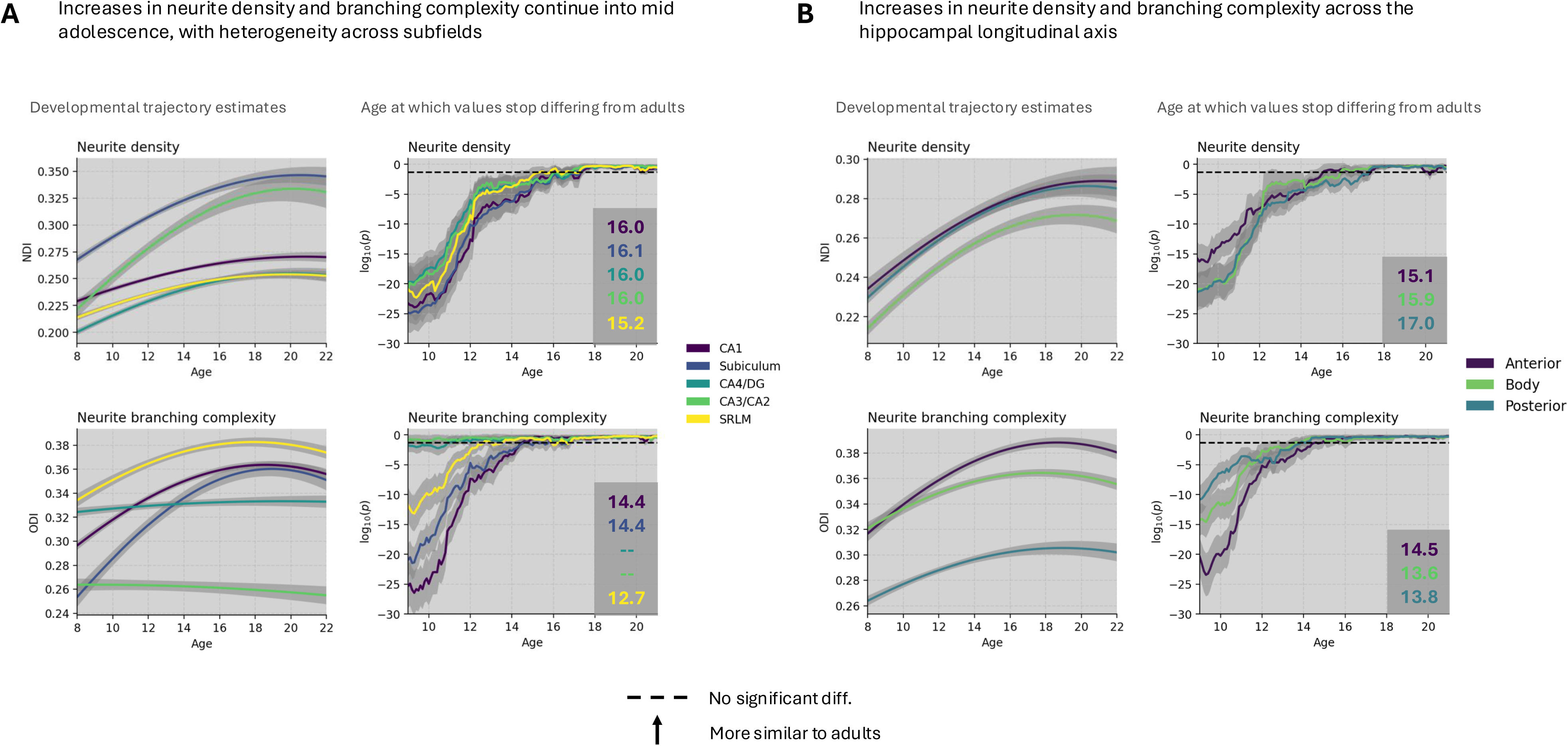
Neurite properties across the hippocampus become adult like by mid-adolescence, with regional heterogeneity. (A) On the left, the nonlinear trajectories overlaid on the same plot. On the right, the log-transformed *p*-values of the *t*-tests comparing subjects at age ± 1 year to adults (higher values reflect greater similarity to adults; dashed-lines reflect no significant difference with adults at *p* = .05). For each subfield, the age at which subjects no longer show a significant difference with adults is presented in the bottom right hand corner. (B) Same as *A*, for longitudinal positions.

Taken together, these results demonstrate that neurite density continues to increase into mid-adolescent development across the entire hippocampus, with developmental effects that are longer-lasting in the posterior relative to anterior hippocampus, yet relatively consistent across subfields. Neurite branching complexity, on the other hand, matures at an earlier age and exhibits more heterogeneity across the hippocampus, with developmental effects that are longer-lasting in anterior relative to posterior hippocampus, as well as in CA1, SRLM and subiculum relative to CA4/DG and CA3/CA2, which were already adult-like in the younger children in our sample.

### Age-related differences in neurite properties spatially correlate with the cell-type composition of hippocampal tissue

Our results so far show non-linear increases in both neurite density and branching complexity across the hippocampus into mid-adolescence, with developmental effects varying by subfield and along the longitudinal axis. However, with these analyses alone, we are unable to tease apart the cellular composition of hippocampal tissue that exhibits these developmental changes in neurite (e.g., are increases in neurite density primarily attributable to changes in glia or pyramidal neurons?). To address this question, we devised a novel technique in which we integrate our imaging data with transcriptomic data from the Allen human brain atlas, as well as hippocampal cell-type gene-expression signatures identified via single-nuclei RNA sequencing (Ayhan et al., 2021). With this technique in hand, we then characterize the cellular composition of hippocampal tissue which exhibits developmental change in neurite density and branching complexity.

An outline of this technique is shown in Fig. 5a. First, we extracted gene-expression data for 107 MNI-coordinates of the bilateral hippocampus using the Allen human brain gene-expression atlas (AHBA; Hawrylycz et al., 2012). Then, at each of these locations, we used the gene-expression data to estimate the signatures of 34 cell-types that exist within the adult hippocampus (Ayhan et al., 2021). These cell-types, which include; 6 pyramidal neuron subtypes, 5 inhibitory neuron subtypes, 7 granule neuron subtypes, and 16 glial cell subtypes, are found across the whole hippocampus, with prominent variation across both the longitudinal axis and subfields (Ayhan et al., 2021). The output of this analysis step was a [107-region by 34-cell-type] matrix quantifying the gene-expression signature of each cell-type at each hippocampal location in the AHBA (bottom panel of Fig. 5a) Many cell-type gene-expression signatures are highly correlated with one another (i.e., the signatures of all granule and inhibitory cell subtypes are highest at dentate gyrus locations). To account for these correlations in cell-type signatures, and facilitate integration with our imaging data, we then decomposed this [107 by 34] matrix into a [107 by 6] matrix using non-negative matrix factorization (Fig. 5b). This analysis provided us with 6 spatial components, each of which is related with a specific mixture of hippocampal cell-types. The cell-type loadings of these components, grouped by [pyramidal, inhibitory, granule, glial] are shown in Fig. 5b and in more detail in Supplementary Fig. 5. As can be seen from inspection of the NMF component matrix, the major patterns of cell-type prominence were captured, with NMF_1 exhibiting high loadings of granule cells and inhibitory neurons in DG, and NMF_2 exhibiting high loading of pyramidal neurons in CA1.

**Fig 5.**
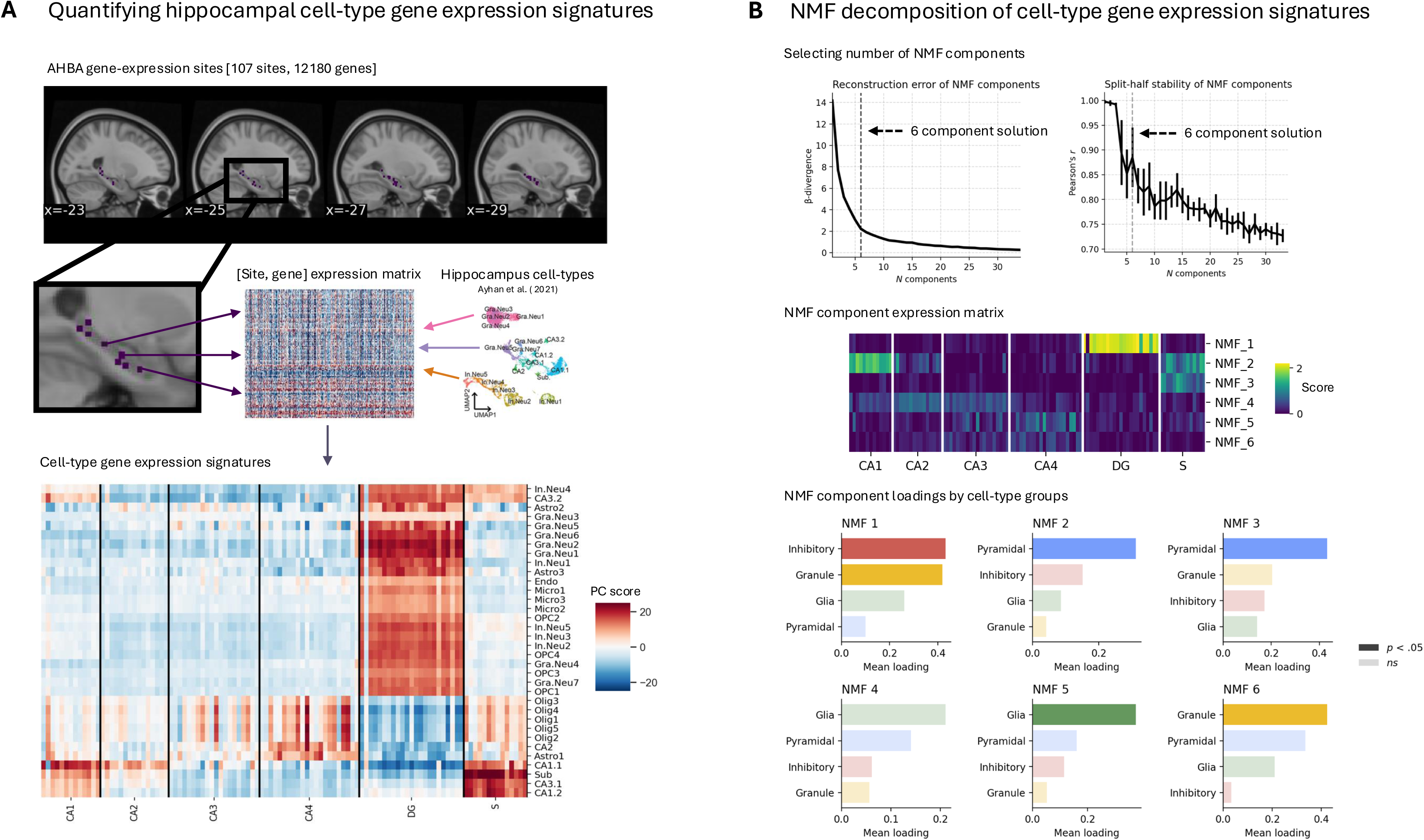
Pipeline for quantifying the cell-type composition of hippocampal tissue. (A) Top panel: sagittal cross-section of an MRI showing the 107 hippocampal locations in the Allen Human Brain atlas (AHBA) used in analyses. Bottom panel: the cell-type expression signatures calculated from the gene-expression data. Note: Insert of the UMAP cell types is adapted from Ayhan et al. (2021), Figure 2. (B) Top panel: reconstruction error and split-half stability of the NMF decomposition as a function of the number of components. The 6-component solution, capturing the majority of variance in the cell-type expression signature matrix with high stability, is highlighted with the dashed line. Middle panel: 6 NMF components used in subsequent analyses. Bottom panel: loadings of cell-type groups for each component. Cell-type groups (e.g., pyramidal neurons) that are over-represented on a given component (*p* < .05) are darkly shaded.

Once the hippocampal cell-type matrix had been decomposed into 6 spatial components, we correlated the scores of these components with age-related changes in neurite density [NDI] and branching complexity [ODI] across the 107 locations in the hippocampus. Age-related changes in neurite density at each of the 107 hippocampal sites were quantified as the correlation between age and NDI/ODI. We found a significant positive correlation between age-related changes in neurite density and the spatial component NMF_6 (*r* = .23, *p* = .03; Fig. 6). The cell-type composition of this component, along with its distribution across the hippocampus, can be seen in Fig. 6. This component was significantly over-representative of granule cell-type signatures (primarily Gra.Neu5, Gra.Neu3, Gra.Neu6, and Gra.Neu4) as well as Astro1 and CA1 cells, and was most prominent at CA_4 and CA3 sites. It did not differ significantly along the longitudinal axis (Fig 6A, bottom right). We also found a significant positive correlation between age-related changes in neurite branching complexity and the spatial component NMF_2 (*r* = .25, *p* = .03; Fig. 6). The cell-type composition of this component, along with its distribution across the hippocampus, can be seen in Fig. 7. This component was significantly over-representative of pyramidal cell-type signatures (primarily CA1.1, CA3.2, Sub, CA1.2 and CA2), was most prominent at CA1 and subiculum sites, and correlated significantly with position along the longitudinal axis, increasing towards anterior hippocampus.

**Fig 6.**
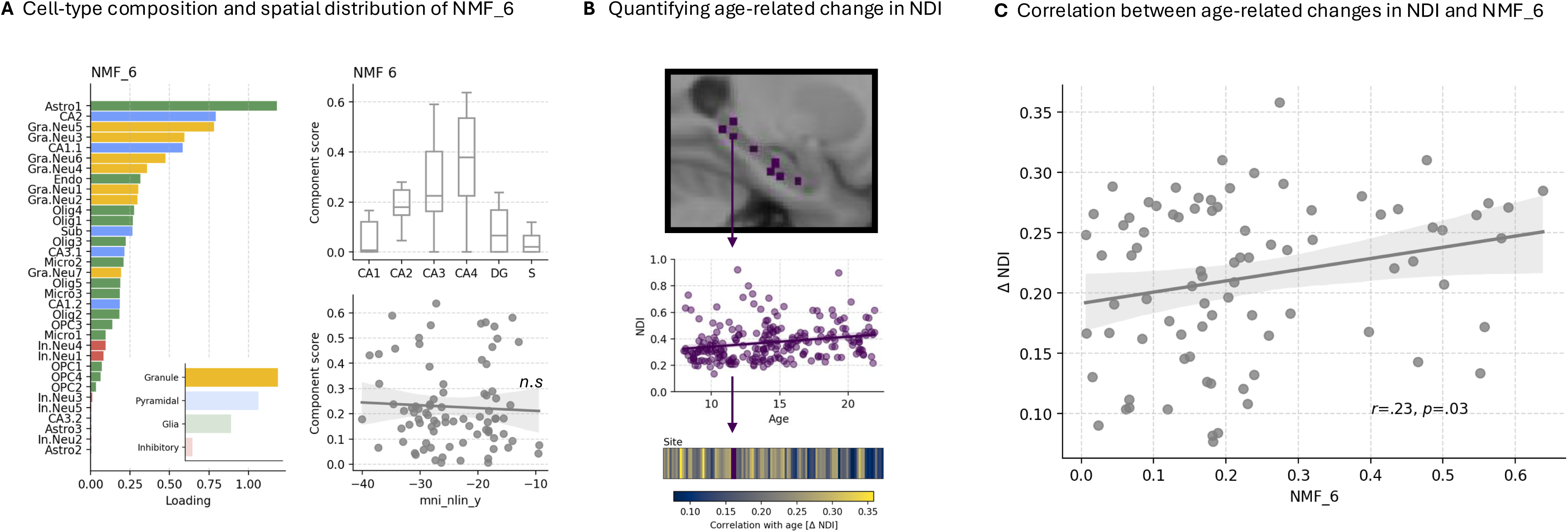
Age-related changes in neurite density overlap with a granule-neuron component. (A) Loadings of the NMF_6 component, color-coded by cell-type grouping (left), along with a boxplot and scatter-plot showing the spatial distribution of NMF_6 across hippocampal subfields (right, top) and the hippocampal long-axis (right, bottom), respectively. Mni_nlin_y: MNI y coordinates (B) Schematic showing how age-related changes in neurite density were quantified: as the correlation between age and NDI at each of the 107 hippocampal sites. (C) Scatterplot showing the correlation between age-related change in NDI and the NMF_6 component.

**Fig 7.**
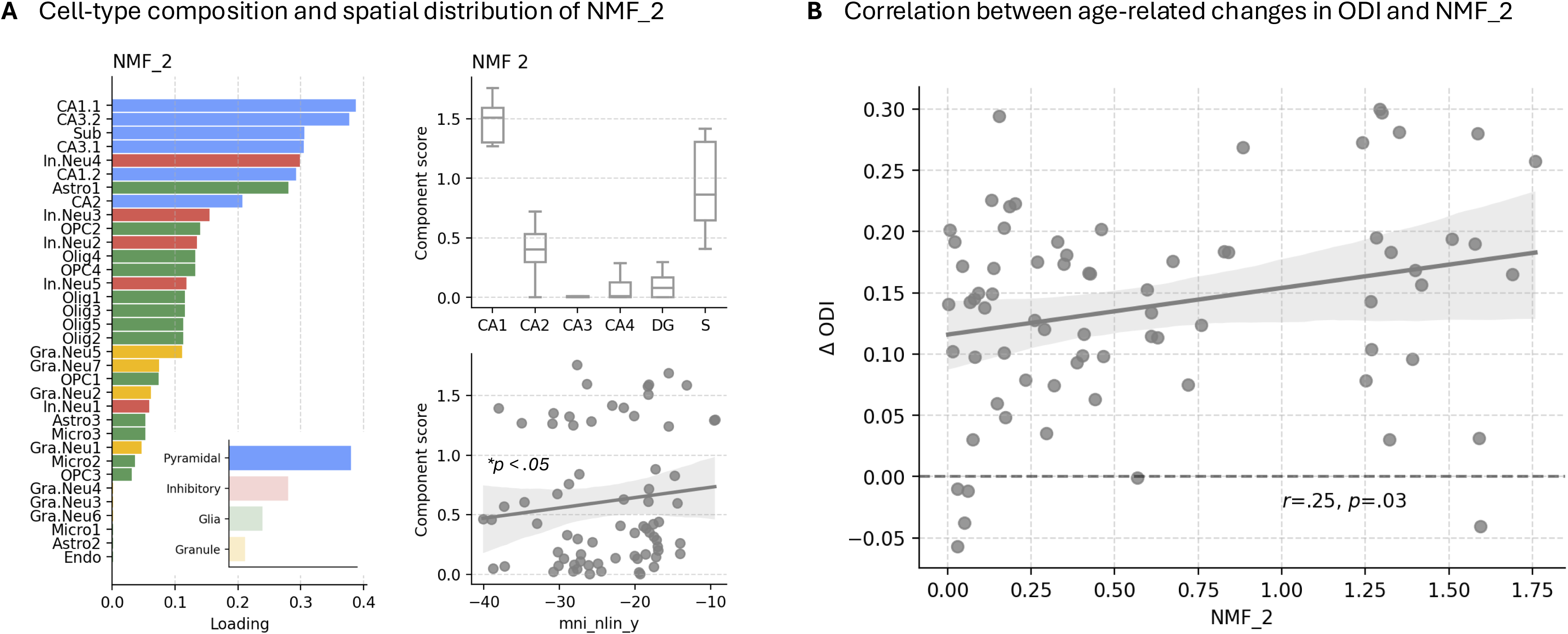
Age-related changes in neurite branching complexity overlap with a pyramidal-neuron component. (A) Loadings of the NMF_2 component, color-coded by cell-type grouping, along with a boxplot and scatter-plot showing the spatial distribution of NMF_2 across hippocampal subfields and the hippocampal long-axis, respectively. (B) Scatterplot showing the correlation between age-related change in ODI and the NMF_2 component.

### Imaging markers of neurite development are spatially correlated with genes related to neuronal and glial projections and their development

We also found a colocalization between age-related change in neurite and the expression of specific genes: 993 genes overlapped with age-related change in neurite density, and 220 genes overlapped with age-related change in neurite branching complexity (Pearson’s *r, p* < .05). The functional profiles of these gene sets were then interrogated through a gene-ontology analysis. We found that the 993 genes which overlapped with age-related change in neurite density were significantly enriched for 39 cellular components and 33 biological processes (FDR-*p* < .05), all of which can be seen in Figure 8. No significant terms were associated with the 220 genes that spatially overlapped with age-related change in neurite branching complexity.

**Fig 8.**
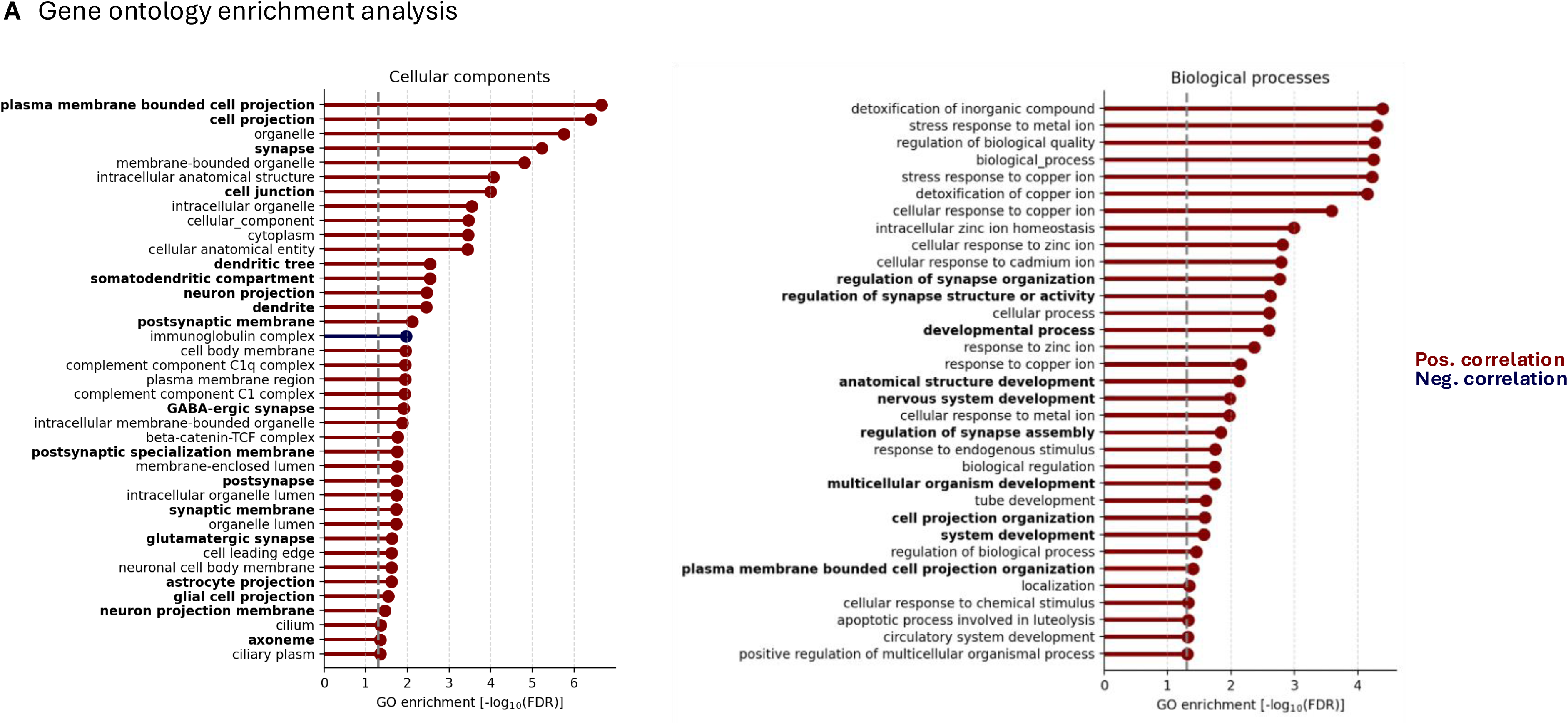
Gene-ontology analyses. Genes significantly correlated with age-related change in neurite density (FDR-*p* < .05) were subjected to a gene ontology analysis. Shown are the cellular components and biological processes associated with this gene set (at FDR-*p* < .05; gray dashed-line). Larger GO enrichment values indicate greater enrichment.

Inspection of the cellular components revealed an association with neurites, including dendrites and glial processes (example terms: dendrite, neuron projection, glial cell projection, cell projection, dendrite, and astrocyte projection). Inspection of the biological processes revealed an association with neurodevelopment in general (e.g., ‘nervous system development’), and neurite development in particular (e.g., ‘regulation of synapse assembly/synaptogenesis’).

## Discussion

Using a biophysical model of diffusion-weighted MRI sensitive to the density and branching complexity of neurites, we offer the first glance at how these properties come to vary along the prominent axes of hippocampal organization throughout childhood and adolescence. We found the density and branching complexity of neurite both continue to increase until mid adolescence, approximately 14 to 16 years of age, at which point they reach adult-like values. This relatively protracted maturation is similar in timing to the development of neurite density and branching complexity observed across association areas in the frontal cortices (Lynch et al., 2023). While increases in the density of neurite were observed across the entire hippocampus (with prominence in the posterior relative to anterior portion of the hippocampus), increases in the branching complexity of neurite were heterogeneous across the hippocampus, primarily confined to CA1/SRLM, the subiculum, and the anterior portion of the hippocampus.

By relating these results with an atlas describing the cell-type composition of hippocampal tissue, we found that increases in neurite density overlapped with a spatial component composed primarily of CA4 granule neurons. Granule neurons synapse onto pyramidal neurons in CA3, forming the mossy fibre connections of the trisynaptic hippocampal circuit (Neves et al., 2008). Our results may therefore reflect maturation of this mossy fiber pathway. We also found that increases in neurite branching complexity overlapped with a spatial component composed primarily of pyramidal neurons, particularly those within CA1. Our measure of branching complexity, the orientation dispersion index of the NODDI model, is thought to be particularly sensitive to basal dendrites. This is because basal dendrites typically promote molecular diffusion perpendicular to the principal direction of diffusion, which is most often aligned with apical dendrites (Wang et al., 2019). With this in mind, the increases we observed in ODI may reflect an increased branching of the basal dendrites of CA1 neurons.

Our results are strengthened by several factors. First, the relatively homogenous increases in neurite density we observed across the hippocampus are in line with previous work from Mah et al. (2017), who found NDI across the whole hippocampus to increase with age from 8 to 13 years old. While Mah et al. (2017) did not observe any age-related differences in neurite branching complexity, it is likely in light of current findings that averaging across the entire hippocampus obscured the age effects, since changes in branching complexity varied considerably across subregions. Second, the relative differences in NDI and ODI we observed across subfields and the longitudinal axis (particularly in the adults of our sample; see supplementary Fig. 2) are highly consistent with previous findings in adults (Karat et al., 2023). Specifically, the density of neurite is lower in CA1 and CA4/DG when compared to CA3, CA2, and subiculum; and the branching complexity of neurite is highest in CA1, subiculum, and anterior relative to posterior hippocampus (Karat et al., 2023). Finally, our results are strengthened by the large sample size (*N*=364), high-resolution T1 images (submillimeter), and accurate segmentation protocol (MAGeT; Chakravarty et al., 2013).

Despite these strengths, a limitation of the current work is that our gene-expression data and thus cell-type expression signatures were acquired from an adult sample. Given known changes in gene-expression throughout childhood and adolescence, it is unclear the extent to which the spatial patterning of our cell-type expression signatures are valid for cross-reference with children. However, we have reasons to believe that these cell-type expression maps are interpretable. First, we found that the genes which are spatially correlated with age-related changes in neurite density are over-represented in cellular components and biological processes related to neurite development (terms include: dendrite, dendritic tree, neuron projection, glial cell projection, astrocyte projection, somatodendritic compartment, nervous system development, regulation of synapse organization; see supplementary Figure 6). These results suggest there is likely a spatial alignment of our data with the adult-derived gene-expression data. Second, the cell-type composition of tissue across the hippocampus primarily differs according to subfield (Ayhan et al., 2021), and the cell-type differentiation of hippocampal subfields primarily occurs much earlier in development: prenatally during the third trimester (Lee et al., 2017).

These results provide the first description of how neurites in the human hippocampus mature throughout child and adolescent development: in a subregion-specific way that aligns with the cell-type composition of hippocampal tissue and reaches maturity by approximately mid-adolescence. This maturation was closely aligned with functionally-relevant axes of hippocampal organization, suggesting these changes may be associated with the changes in hippocampal function observed throughout this period (e.g., Calabro et al., 2020). Understanding this aspect of human neurodevelopment will be important in explaining the increases in memory capacity that are observed throughout this developmental period (Fjell et al., 2019).

## Methods

### Participants

Data for the current study were obtained from the development cohort of the Human Connectome Project (HCP-D; Somerville et al., 2018). Our group obtained a NIMH data archive data use certification from McGill university. Informed consent was obtained during data collection. Subjects in the HCP-D are healthy/typically developing; i.e., they were subjected to extensive exclusion criteria including: prior head injuries, psychiatric illness (Schizophrenia, Bipolar disorder, Anxiety, Major depression), and the reception of special services at school (e.g., for dyslexia, intellectual disabilities, speech/language pathology; see full criteria in Somerville et al., 2018). All unrelated subjects were initially included in analyses (*N* = 545). Two stringent quality-control procedures were applied (described in greater detail below), resulting in the removal of 136 subjects due to low quality T1 images, and removal of 45 subjects due to low-quality alignment of diffusion-weighted and T1 images (≥ 10 ‘misaligned’ voxels, classified as non-gray matter prior to transformation and within the hippocampus following transformation). This resulted in a total of 364 subjects entering analyses, aged 8.0 to 21.9 years old (*M* = 14.14, *SD* = 3.99), with 193 females and 171 males.

### MRI data acquisition

Images were acquired with a Siemens 3T Prisma scanner (80 mT/m gradients, slew rate of 200T/m/s) and Siemens 32-channel Prisma head coil (Harms et al., 2018). One T1-weighted image and 21 minutes of diffusion-weighted imaging were acquired (Harms et al., 2018). T1-weighted images had 0.8 mm isotropic voxels (sagittal FOV = 256 x 240 x 166 mm, matrix size = 320 x 300 x 208 slices, slice oversampling of 7.7%, 2-fold in-place acceleration in phase encode direction, bandwidth = 744 Hz/pixel), TR/TI = 2500/1000, TE = 1.8/3.6/5.4/7.2 ms, a flip angle = 8°, and up to 30 TRs. To suppress signals from fat (bone marrow and scalp fat), water excitation was employed. Diffusion weighted images (DWI) had an MB factor of 4, 1.5 mm isotropic voxels, TR = 3.23 s, partial Fourier factor = 6/8, and no in-plane acceleration. 185 diffusion-weighting directions, each with opposite phase encoding (AP and PA) on two shells (*b* = 1500 s/mm^2^, 3000 s/mm^2^), were used over four dMRI runs with 28 b = 0 s/mm^2^ volumes interspersed (Harms et al., 2018; Somerville et al., 2018).

### Hippocampus segmentation

Raw T1-weighted images were preprocessed using the minc-bpipe-library pipeline [https://github.com/CoBrALab/minc-bpipe-library], which includes N4 bias field correction, registration to MNI space (ICBM 2009a Nonlinear Symmetric), brain extraction using BEaST (Bussy et al., 2021), and standardization through ANTs, minc-toolkit, and MNI priors. The hippocampus was segmented into CA1, subiculum, CA4/dentate gyrus, CA2/CA3, and stratum radiatum/lacunosum/moleculare (SRLM) using the Multiple Automatically Generated Templates (MAGeT) technique (Chakravarty et al., 2013). Segmentations were run separately for left and right hemispheres, and manually controlled for quality, resulting in 409 unique subjects (327 with right hemisphere data; 268 with left hemisphere data).

### Diffusion-weighted image processing

Raw diffusion weighted images were denoised using Marchenko-Pastur PCA (MRtrix3; Tournier et al., 2019), corrected for eddy current-induced distortions and subject head motion (FSL; Jenkinson et al., 2012), and corrected for distortion due to magnetic field nonlinearity (FreeSurfer; Fischl et al., 2012). The NODDI model was fitted with the parameters specified in the *WatsonSHStickTortIsoV_B0* model (those originally described in Zhang et al., 2012) with the exception of intrinsic parallel diffusivity (*d*_||_), which was fixed at 1.10 [m^2^/s. Contemporary evidence suggests this may be the optimal choice when interrogating gray matter (Guerrero et al., 2019; Kraguljac et al., 2022; Genç et al., 2018; Fukutomi et al., 2018; Karat et al., 2023).

NODDI was fit in native space, and transformations registering NODDI parameter volumes to population space were created and applied using Advanced Normalization Tools (*ants*; Avants et al., 2011). For each subject, all *b* = 0 diffusion-weighted images were averaged, and this mean *b*0 image was rigid-transformed to the T1 image. Then, the T1 image (which had the highest resolution) was non-linearly registered to the population average. This transformation was created separately for right and left hemispheres, and was focused with a ‘bounding-box’ inclusive of the hippocampus. This bounding box was created in population space, and included any voxels that fell within the hippocampus of a single subject, plus 5mm along each *x, y,* and *z* direction. To ensure accurate transformations, T1 images were segmented into 3-tissues (WM, GM, CSF) using FSL’s *fast*, and subjects with transformations that resulted in 10 or more ‘misaligned’ voxels (i.e, voxels that fell within the hippocampus yet were not classified as gray matter) were removed from further analyses, leaving 279 subjects with right hemisphere data and 230 with left hemisphere data. Segmentations were then manually inspected by author JK, although this did not result in the removal of any subjects. The voxel-wise group-average values of NODDI parameters (NDI and ODI) are shown in supplementary Fig. 3.

#### Nonlinear developmental trajectories

To identify non-linear changes in age, we fit the *KernelRidge()* transformer in *scikit-learn* with the radial basis function kernel. The width of this kernel (□-parameter), controls the amount of smoothing (i.e., a sufficiently large kernel will converge to a linear fit). This parameter was selected separately for each metric-subregion pair through 5-fold cross-validation so as to maximize the out-of-sample *R^2^*. The 95% confidence intervals were defined as ± 1.96 times the bootstrapped standard error of the mean, derived from 100 bootstrapped samples.

To identify the age at which NDI and ODI values stopped differing significantly from adults, we conducted running t-test analyses with a sliding time-window. Specifically, at each age (0.1 year increments), we conducted an independent samples *t*-test between all subjects that were aged ± 1 year from the selected age (resulting in a 2-year window) and the adults in our sample (18 years or older; *N*=80). We then took the *p*-values from these tests, and identified the minimum age at which these *p*-values were greater than 0.05. The standard error was estimated through 100 bootstrapped samples of the data.

### Imaging transcriptomics

#### Preprocessing

Data were cross-referenced against post-mortem gene expression data from the Allen Human Brain Atlas (AHBA; Hawrylycz et al., 2012). To increase reproducibility, we obtained a subset of AHBA data that had already been processed for hippocampus-specific analyses by Vogel et al. (2020). Probe values in this preprocessed data are the standardized residuals from a linear model that included dummy-coded donor-ID variables; thus, probe values reflect average gene-expression, independent of inter-individual differences. MNI coordinates in this data are those re-processed by Devenyi (2018) to account for errors that arose from transformations into MNI space (Arnatkevic□iūtė et al., 2019, Vogel et al., 2020).

From this data, genes not expressed in the hippocampus (according to the Human Protein Atlas: [https://www.proteinatlas.org]) were filtered out. Then, since multiple probes index the expression of single genes in the AHBA, we took the mean expression across probes for each gene (Arnatkevic□iūtė et al., 2019). To exclude sites whose voxels did not overlap with our subject’s hippocampi, we first created right and left ‘majority-vote masks’, which included voxels that fell within the hippocampus of half our subjects or more. Then, gene-expression sites whose MNI coordinates fell outside the hippocampal mask were removed. Finally, for a given gene-expression site, the MNI coordinate voxel, and all 26 of its immediately adjacent voxels, were defined as a neighbourhood, resulting in a 27mm^3^ box for each gene-expression site. The output of these analyses was a 107-region by 12,180-gene matrix with normalized gene expression levels for all hippocampal sites. To quantify the age-related change in NDI and ODI at a given gene-expression site, we took the correlation of a given metric with age (Pearson’s *r*; see Fig. 6). Gene ontology analyses were performed at [http://geneontology.org/] for Homo Sapiens.

#### Cell-type expression

To quantify cell-type gene-expression signatures, we obtained the cell-type gene sets from Ayhan et al. (2021). These cell-types were acquired from single-cell RNA sequencing of the adult human hippocampus. Then, we took the weighted expression of genes, with weights determined through principal component analysis [PC1 of the cell-type gene-expression matrix], as in prior work (Seidlitz et al., 2020; Vogel et al., 2021). This analysis was identical to the cell-type analyses conducted by Vogel et al. (2021), except we made use of cell-types from more recent work that were specifically identified from hippocampal tissue (as opposed to tissue from the middle temporal gyri). We calculated the gene-expression signature for the 34 hippocampal cell-types identified by Ayhan et al. (2021), which included all neuronal sub-classes, resulting in a [107-region by 34-cell-type] matrix. We then decomposed this into a [107-region by 6-component] matrix using non-negative matrix factorization (NMF). NMF is a dimensionality-reduction technique which decomposes sample-by-feature matrices into components with sparse and non-negative loadings; important factors when considering the interpretability of components (Roads & Love, 2024; Patel et al., 2020; Wang et al., 2012). NMF was implemented in *scikit-learn* with the *MinMaxScaler()* and *NMF()* transformers. To select the number of components (*k*), we inspected the reconstruction error and split-half stability across an increasing number of *k*-values. We observed diminishing returns in reconstruction ability following approximately 6 NMF components, suggesting 6 patterns of spatial variation in cell-type expression signatures are adequate to capture the majority of variation in our data (Fig. 5). This 6 component solution also exhibited a relative peak in split-half stability: when NMF was fit on randomly split halves of the data, NMF-scores were highly correlated with one another (*r* = .88), whereas a relative drop-off in stability was observed for higher values of *k*.

## Data availability

Raw data is freely available through the Human Connectome Project, upon completion of a data-usage agreement: [https://nda.nih.gov/general-query.html?q=query=featured-datasets:HCP%20Aging%20and%20Development]. Preprocessed data (volumes and microstructures for subfields and anterior/body/posterior subregions) is available in the Supplementary material. Note: volumes are in mm^3^.

## Code availability

The code required to run analyses has been published on github: [https://github.com/JonahKember/hippocampal_microstructure].

## Supplementary figure legends

**SFig 1.**
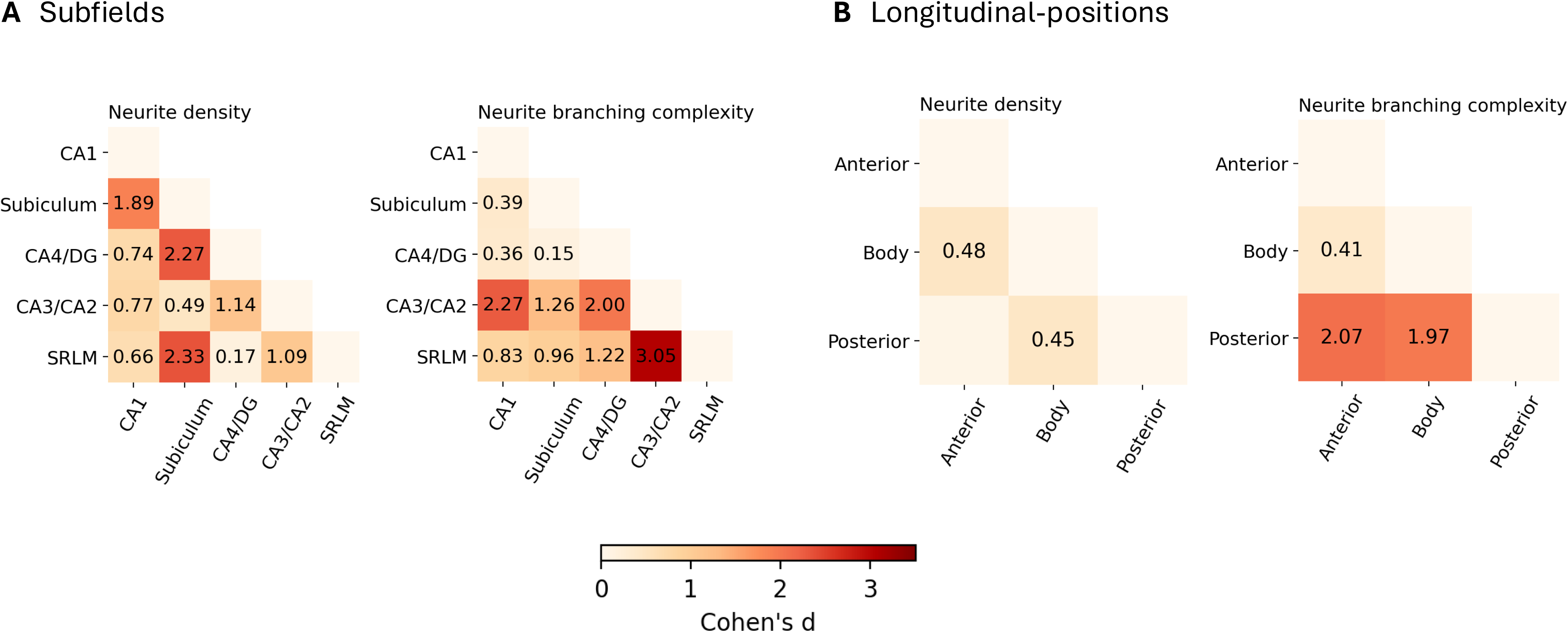
Effect-sizes of differences in neurite density and branching complexity across subfields and longitudinal-positions. (A) Subfield-by-subfield matrices showing the effect-size (Cohen’s *d*) of differences in neurite density and branching complexity. Values not shown are *p* < .05. (B) Same as *A*, for longitudinal-positions.

**SFig 2.**
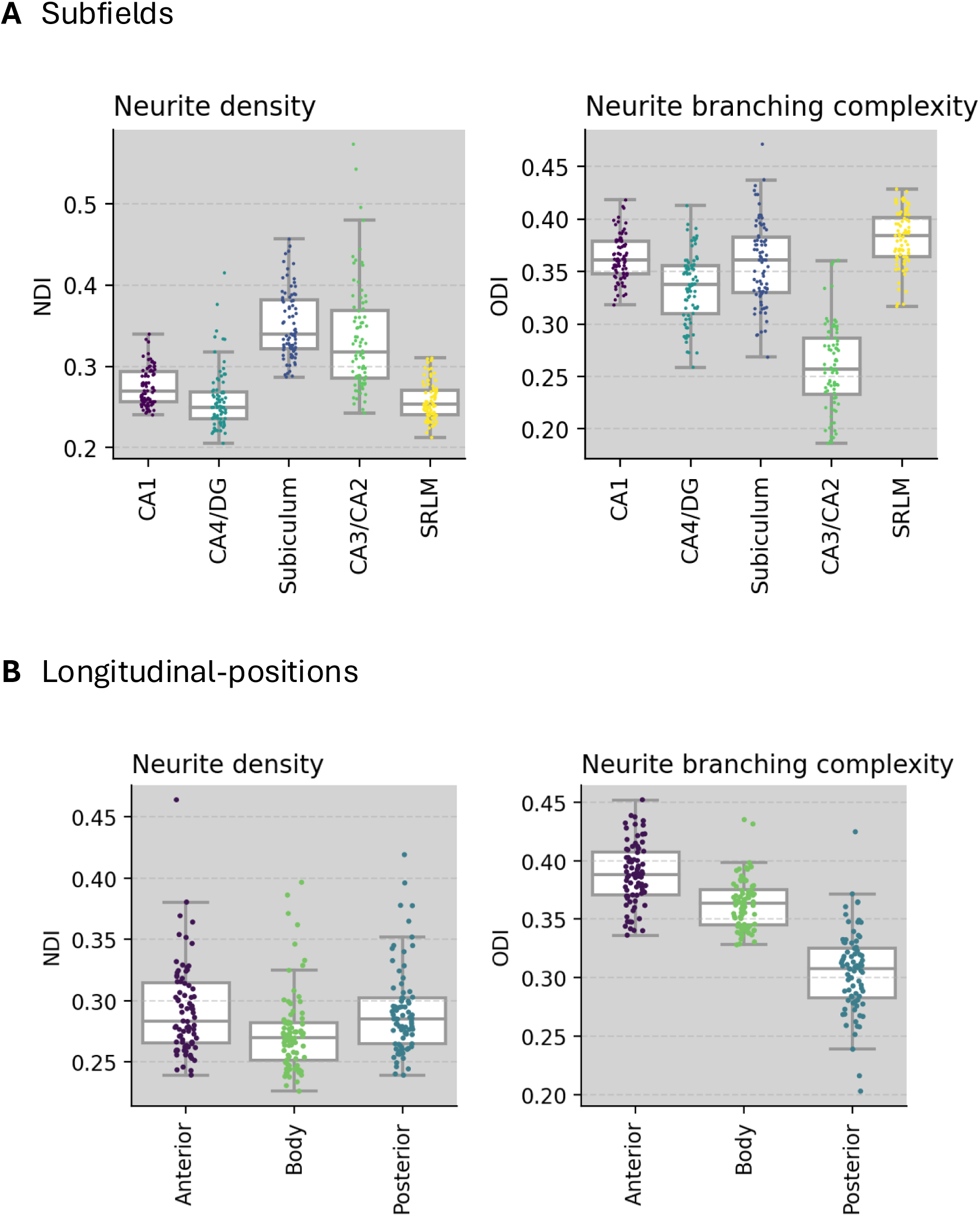
Differences in neurite density and branching complexity across subfields and longitudinal-positions in adults. (A) Box-plots illustrate the average neurite density and branching complexity across hippocampal subfields for each adult subject in our sample (age >= 18; *N*=80). (B) Same as *A*, for longitudinal-positions.

**SFig 3.**
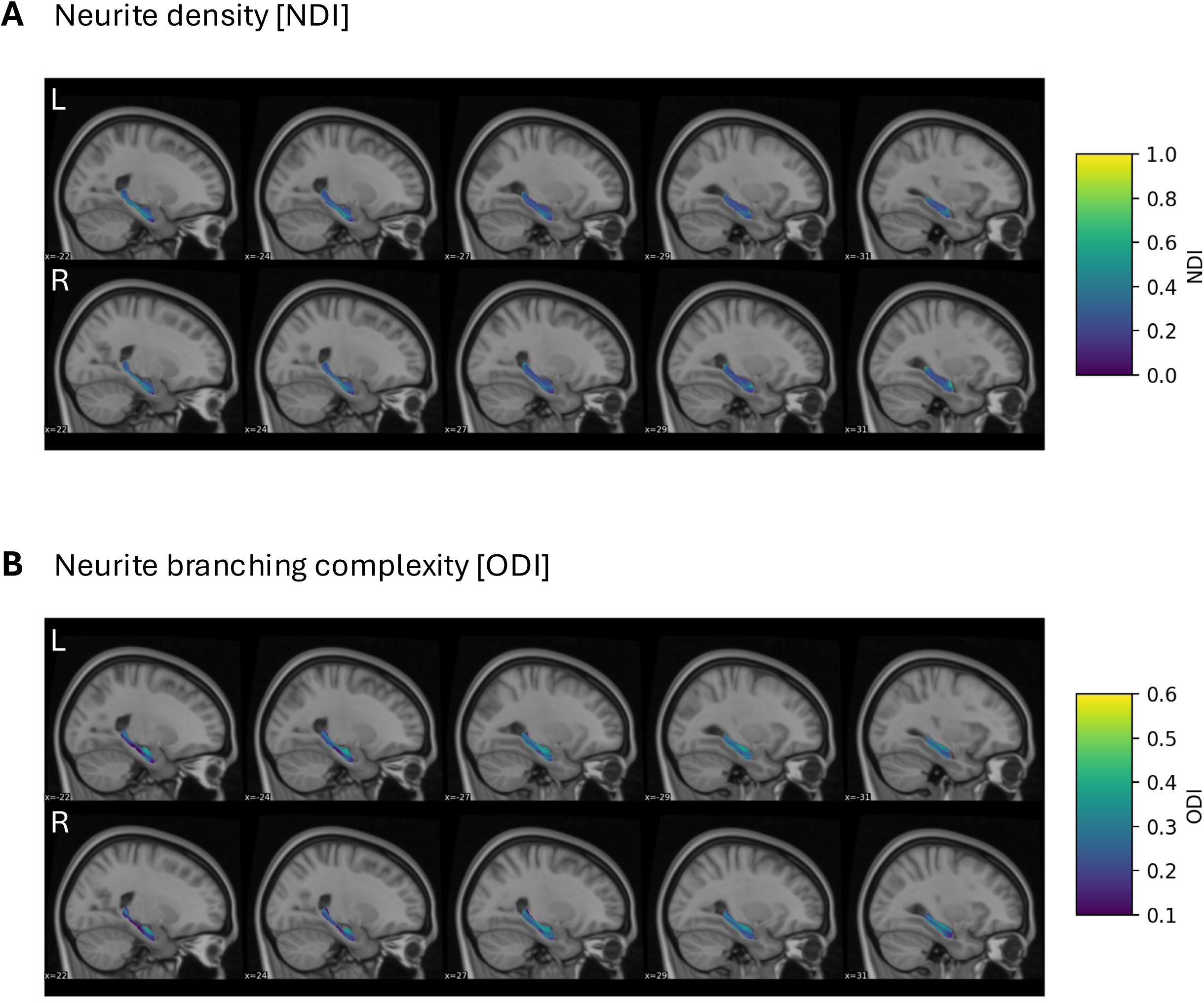
Group-average of voxel-wise neurite density and neurite branching complexity estimates. (A) Sagittal MRI slices showing variation in neurite density across the hippocampus. (B) Same as in *A*, for neurite branching complexity.

**SFig 4.**
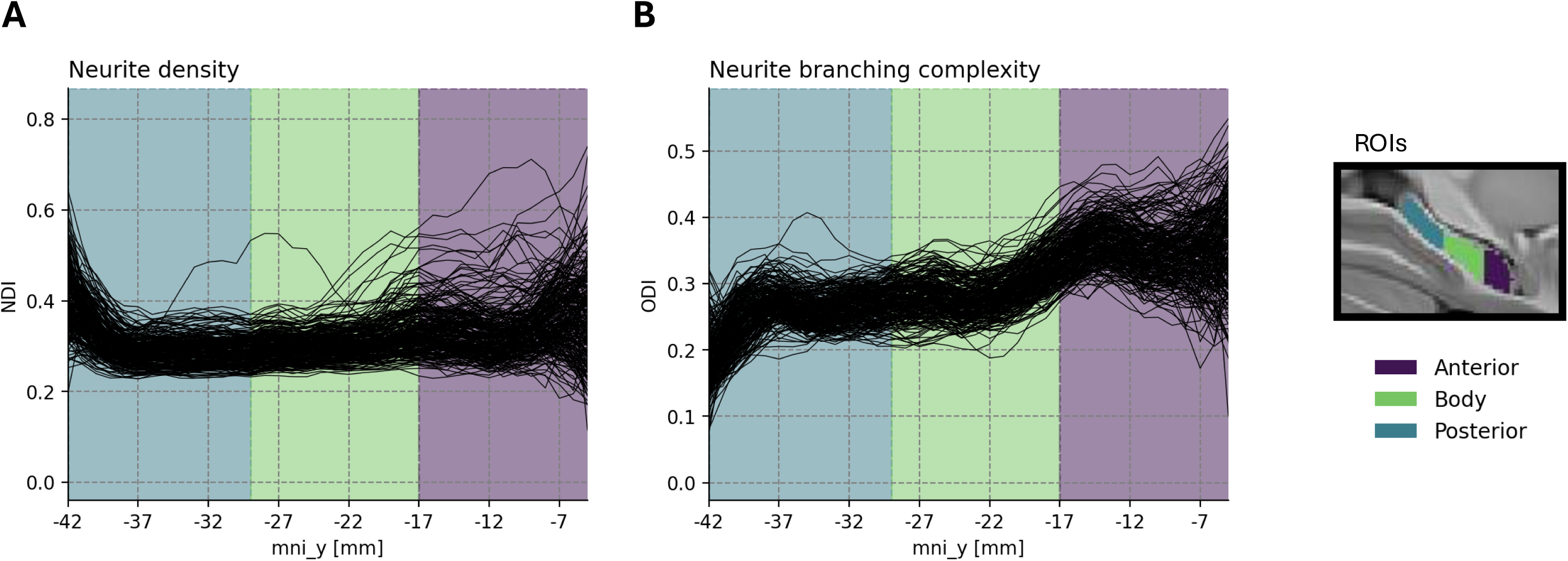
Differences in neurite properties across the *y-axis*. To assess whether results related to the longitudinal axis would be robust to the specific hippocampal parcellation used, we plotted neurite density and branching complexity at each mm of the *y-axis*. As can be seen, the trend of higher ODI values in anterior relative to posterior hippocampus (shown in Fig. 2 of the main text) is visible when inspecting the entire longitudinal axis.

**SFig 5.**
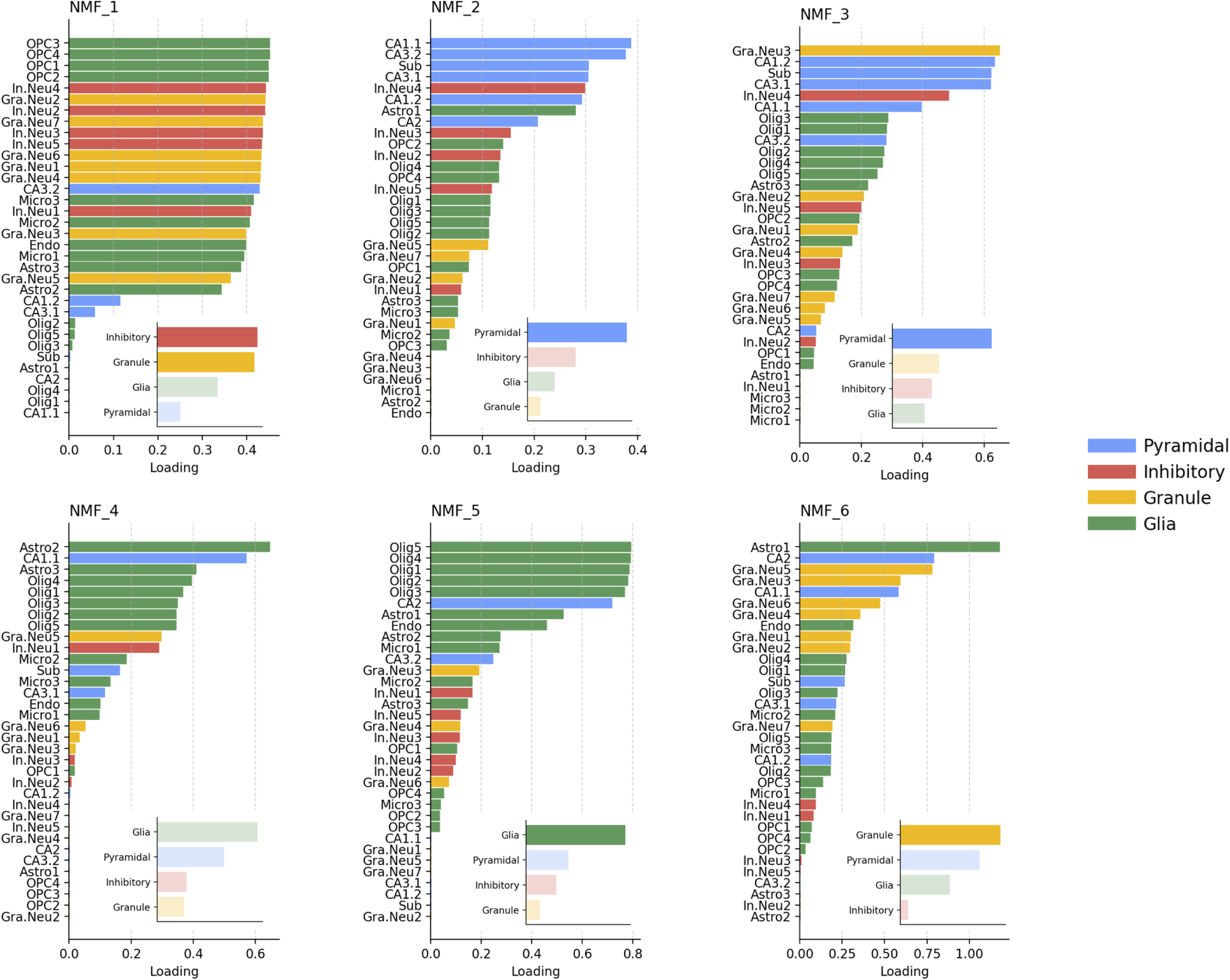
Cell-type loadings for all 6 NMF components. Bar plots showing the loadings of each cell-type onto each NMF component. In the bottom right hand corner of each plot, the average weighting of each cell-type grouping, transparency encodes whether the mean weighting of a given cell-type grouping is significantly greater than expected by chance (shaded = *p* < .05).

